# Development of Advanced Bioinformatic Profiles to Improve the Detection and Functional Understanding of Fungal Acid Phosphatases

**DOI:** 10.1101/2025.08.21.671465

**Authors:** Támara Gomez-Gallego, Zulema Udaondo, Rocio Palacios-Ferrer, Luis Díaz-Martínez, Juan L. Ramos

## Abstract

We have retrieved approximately 9,000 protein sequences annotated as fungal acid phosphatase or phytase from the UniProtKB database. After stringent quality filtering, a curated dataset comprising 3,058 high-confidence sequences was assembled. Phylogenetic analysis revealed that these enzymes segregate into eight distinct clades, representing distinct groups of fungal acid phosphatases, purple acid phosphatases, phytases and groups containing both phytases and acid phosphatases annotations. Based on this classification, we have developed three representative protein profiles referred to as Pfr-A-Fungal_phos, Pfr-B-Fungal_phos and Pfr-C-Fungal_phos, each designed to capture the phylogenetic and functional of the enzyme families. The specificity and breadth of these profiles were validated through comprehensive heat-map analyses. When deployed on public protein and metagenomic databases, these profiles demonstrated high sensitivity and specificity, enabling the identification of hundreds of previously uncharacterized fungal proteins. These proteins spanned a broad taxonomic distribution, with notable prevalence in the *Ascomycota* and *Basidiomycota* phyla. Collectively, these findings highlight the utility of the newly developed profiles to uncover a novel and taxonomically widespread family of fungal enzymes. This work provided valuable insights into the evolutionary diversity and ecological significance of fungal enzymes involved in phosphorous cycling.

## INTRODUCTION

Phosphorus (P) is an essential macronutrient required for the synthesis of key cellular components, including nucleic acids, phospholipids, and energy carriers such as pyrophosphate, ADP and ATP, as well as various organophosphorus molecules vital to all living organisms (Malhotra et al., 2018). Although abundant in the Earth’s crust, most P exists in insoluble forms, making it largely unavailable to plants, macroalgae, and microorganisms. The global biogeochemical cycling of P is crucial for ecosystem productivity and is closely linked to carbon and nitrogen cycles (Moore, 2013). Unlike carbon and nitrogen, the phosphorus cycle lacks a significant gaseous phase, relying primarily on rock weathering and, to a lesser extent, dust deposition for input (Turner et al., 2012), rendering P availability a limiting factor in many ecosystems (Walker & Syers, 1976). Phosphorus is also vital for crop productivity, yet its availability in many soils is limited and depends on finite phosphate rock reserves (Cordell et al., 2009).

Microorganisms, particularly bacteria and fungi, play a central role in mobilizing phosphorus from organic and inorganic sources (Brazhnikova et al., 2022). Phosphate-solubilizing microbes (PSMs) convert insoluble phosphorus into bioavailable forms. Inorganic phosphates, such as aluminum, calcium, and iron phosphates, are solubilized through the microbial production of organic acids (e.g., gluconic, oxalic, lactic, malonic), which acidify the local environment and enhance P solubilization. For organic phosphorus compounds, microbes and plants rely on the secretion of phosphatase enzymes that hydrolyze these compounds releasing ortho phosphate. Phosphatases are broadly categorized into two classes based on their optimal pH: i) alkaline phosphatases (EC 3.1.3.1) that function preferentially in the range of pH between 7.5 and 8.5, and are predominantly of microbial origin (Tarafdar & Claassen, 1988). These enzymes, encoded by gene families such as *phoA*, *phoD*, *phoX*, and *psiP1* in bacteria, and *pho8*, *xpp1*, *ssu78*, *siw114*, and *alp1* in fungi, are widespread in soils and marine environments (Ragot et al., 2015; Neal et al., 2018; Duhamel et al., 2025; Jin et al. 2022; Balenga et al.,2015); and ii) Acid phosphatases (EC 3.1.3.2) that function optimally at low pH, i.e. in the range of 5 to 6. These enzymes, commonly released by plants and microbes (Yadav & Tarafdar, 2001), hydrolyze phosphoesters and phosphoanhydrides. Several, subtypes of fungal acid phosphatases have been described based on their cellular localization, their substrate preference and their differences in regulation (Straket and Mitchell, 1986; Mitchell et al., 1997; Fujita et al., 2003). At the molecular level, two main molecular groups of acid phosphatases have been consistently described in fungi that includes, histidine acid phosphatases (HAPs), which includes a phytase sub-group -, and the metallophosphoestarases, purple acid phosphatases (PAPs). HAPs are a broad family that includes enzymes with a conserved catalytic histidine residue and display a RHGXRXP motif. Members of this group of enzymes show a broad substrate specificity and are active across a range of acidic pH values (Mitchell et al., 1997). The phytase subgroup of HAPs (EC 3.1.3.8, EC 3.1.3.26) are specialized in hydrolyzing phytic acid (phytate) (myo-inositol hexakisphosphate), a major organic P reserve in seeds, releasing phosphate in both acidic and alkaline environments (Lim et al., 2007). The PAPs are a distinct group of metallophosphoesterases with a Fe(III)-tysosinate center that gives them a characteristic purple color. This group of fungal acid phosphatases share a conserved active site motif with plant and mammalian PAPs, but lack the RHGXRXP motif found in histidine acid phosphatases, distinguishing them mechanistically from HAPs (Gocheva et al., 2024).

Soil phosphatase levels are influenced by environmental factors such as total nitrogen, precipitation, temperature, and agricultural practices such as reduced tillage (Margalef et al., 2017; Asensio et al., 2024). Phosphatase activity is commonly assessed using the artificial substrate *p*-nitrophenyl phosphate (pNPP), preferentially reflecting the activity of soil-resident microbial communities. These enzymes facilitate organic matter decomposition and improve plant nutrient uptake, highlighting the microbial contribution to soil fertility. In a previous study, we analyzed bacterial acid phosphatases and, using a refined set of 3,741 sequences, identified three main groups—B, C, and Generic Acid Phosphatases (GAP, Group A) (Udaondo et al., 2020). Phylogenetic analysis revealed the evolutionary relationships of the three groups of bacterial phosphatases, with groups A and C being more closely related than group B. The constructed sequence profiles, specific for each group enabled the identification of many previously uncharacterized acid phosphatases.

Given the high abundance in soil of both phosphate solubilizing bacteria and phosphorous solubilizing fungi, we have now examined fungal acid phosphatases and constructed a phylogenetic tree that revealed eight distinct groups within the fungal kingdom. From this, we have developed three selective SwissProt profiles: one specific to Group I, another to Group II, and a third covering Groups III–VIII. These profiles provide a valuable tool for integrating high-throughput sequencing with field data, improving models of phosphorus cycling, and adding information on the design of microbial biofertilizers.

## MATERIALS AND METHODS

### Phylogenetic tree construction

Protein sequences were retrieved from UniprotKB database (https://www.uniprot.org/) using advanced searches, filtering for fungal sequences annotated with either “acid phosphatase” or “phytase” in the protein name. This search yielded 5,269 sequences annotated as “acid phosphatase” and 3,590 sequences annotated as “phytase”, resulting in total of 8,859 protein sequences (data as of May 6, 2023). Sequences were filtered based on length (mean ± 1 standard deviation), and outliers that were significantly longer or shorter were excluded. Redundant sequences with 100% identity were discarded using EMBOSS Skipredundant software v6.6 (Rice et al., 2000). After manual curation, the data set was reduced to 4,004 sequences. These sequences were aligned using MUSCLE v3.7 software with the parameters —maxiters 1,000 (Edgar, 2004). Highly divergent sequences were removed to improve alignment quality. A final set of 3,058 sequences was retained, re-aligned as detailed above, and used to construct a phylogenetic tree using the IQ-TREE software v1.6.12 (Nguyen *et al*., 2015). The following parameters were used for tree construction: -nt AUTO, -bb 1,000 ; -m TESTMERGE; and -safe, allowing exploration of phylogenetic relationship among sequences. The maximum likelihood tree was constructed using the WAG+I+G4 model, identified as the best-fit evolutionary model by ModelFinder (Kalyaanamoorthy *et al*., 2017). The phylogenetic tree was visualized using the Interactive Tree of Life (iTOL) suite, software v4 (Letunic and Bork, 2019).

### Protein profile construction

As an initial step toward creating unique PROSITE generalized profiles, we defined a seed of protein sequences for each enzyme group. The selection of these proteins significantly influences the sensitivity and overall quality of the resulting profiles. For this purpose, full sequences from each group of enzymes were aligned separately using MUSCLE v3.7, as detailed previously. Divergent sequences were further filtered if necessary to enhance alignment consistency. We then used the pfw function from the PFTOOLs v2.3. package to calculate the weights of individual protein sequences within each multiple sequence alignment. The weighted alignments were then converted into generalized profiles using pfmake, applying the parameters: - 2b; -H2.0, with BLOSUM45 as the substitution score matrix. To calibrate the profiles, we used pfsearch and pfscale from the PFTOOLs suite. The calibration database was constructed by filtering fungal proteins from the entire protein Swiss-Prot database (release date Feb 23, 2022), resulting in 34,929 sequences, which were randomly shuffled using a sliding window of 20 residues (-kmer 20) using fasta-shuffle-letters algorithm from MEME suite v5.1.1. After obtaining the calibrated profiles, each profile was compared against the complete database of fungal phosphatases of the phylogenetic tree (3,058 sequences) using pfsearch with parameters –z –f (Sigrist *et al*., 2010), to verify its specificity and ensure it only recognized only sequences from its corresponding group.

### Validation of fungal phosphatase profiles in databases

To further validate protein profiles, each profile was also tested against different protein databases using pfsearch, as detailed above. The resulting match lists were analyzed to warrant satisfactory sensitivity. The database used were: i) the complete dataset of non-curated fungal phosphatase sequences (8,859 protein sequences), ii) fungal protein sequences annotated as “acid phosphatase” or “phytase” in UniProtKB database (20,050 protein sequences, release dated November 21, 2024), iii) the full collection of fungal protein sequences in UniProtKB database (19,274,552 sequences, release date November 21, 2024) and a UniRef90 database containing 29,638,836 clusters of sequences filtered using the following criteria: “Taxonomy: 4751”, which corresponds to Fungi; and “Cluster name: Hypothetical protein OR Unknown protein OR Uncharacterized protein”. The “Cluster” name corresponds to the protein name in UniRef90.

### Cloning and expression of fungal phosphatases in yeast

As a proof of concept to empirically validate fungal phosphatase profiles, the non-revised sequences with the highest Z-score within each profile were synthesized in vitro using codon optimization for *Saccharomyces cerevisiae*.

The selected sequences were: i) a putative purple acid phosphatase from *Hyaloscypha variabilis* (UniProtKB ID A0A2J6S554) for Prf-A-Fungal_phos, herein referred to as HvPhyA,; ii) a putative acid phosphatase/ 3-phytase from *Sclerotinia sclerotiorum* (UniProtKB ID A0A1D9Q6T7) for Prf-B-Fungal_phos, referred to as ScPhyB ; iii) a putative acid phosphatase from *Didymella rabiei* (UniProtKB ID A0A162Z3M2) for Prf-C-Fungal_phos , referred to as DrPhoC. The corresponding DNA sequences were codon-optimized for expression in *S. cerevisiae*, synthesized in vitro (GenScript) and cloned into the yeast expression vector pDR196 (See suppl. Material) between the PstI and Sal I sites. All constructs were verified by sequencing. The BY4741 strain (*MATa his3Δ1 leu2Δ0 met15Δ0 ura3Δ0*) (https://www.yeastgenome.org/strain/by4741, Winston *et al*. 1995; Brachmann *et al*. 1998), was transformed using the lithium acetate-based method (Gietz and Schiestl, 2007) with either the recombinant constructs or an empty pDR196 vector Transformants were selected on synthetic dextrose (SD) minimal medium based on uracil autotrophy. All primers used in this study are listed in Table S1.

### Phosphatase assay in yeast cell cultures

Yeast transformants were grown for 24 h in SD selective medium without uracil. Phosphatase and phytase activities were assayed in cell-free extracts. For this, 20 ml of culture was centrifuged at 10,000 *g* for 10 minutes, the supernatant was discarded, and the pellet was frozen at −80°C. Pellets were resuspended in 3 ml of HAM buffer (40 mM HEPES, 40 mM acetic acid, and 40 mM MES, pH 5.5) (Recio et al., 2024) and cell lysis was achieved using three passes through a French press at a pressure at 1000 p.s.i. The lysates were centrifuged as above and the resulting pellet was resuspended in 1000 µL of HAM buffer .

Acid phosphatase activity was measured by quantifying the amount of *p*-nitrophenol (PNP) produced from 4-nitrophenyl phosphate (PNPP). Briefly, 100 µL of resuspended pellet or supernatant were mixed with 20 µL of pNPP (0.115 M) and incubated for 15 minutes at 37°C. The reaction was stopped by adding 100 µL of NaOH and 0,8 mL of distilled water (Recio et al., 2024). The amount of PNP produced was measured spectrophotometrically at 405 nm using a Tecan Sunrise plate reader (Tecan Austria GmbH).

### Statistical analyses and heatmap representation

All the statistical analyses were performed using the sequence similarities, pairwise sequence similarities were calculated using global pairwise alignment with the align.globalxx function from the Bio.pairwise2 module, which computes percent similarity based on the number of identical matches in the aligned region. The heatmap was constructed using the ComplexHeatmap package (Gu et al., 2016), with color gradients representing similarity values from 0% (blue) to 100% (red) via a midpoint of 50% (white), scaled using colorRamp2 from the circlize package (Gu et al., 2014). Group-level annotations were added as color-coded sidebars using a categorical color palette from RColorBrewer enabling easy visual discrimination among groups.

## RESULTS AND DISCUSSION

We identified eight main groups of protein sequences based on the constructed phylogenetic tree (Fig. 1), and further characterized them using information from the UniProtKB database. A summary of the sequence details is presented in Table 1, with extended raw data available in Table S2.

**Figure 1:**
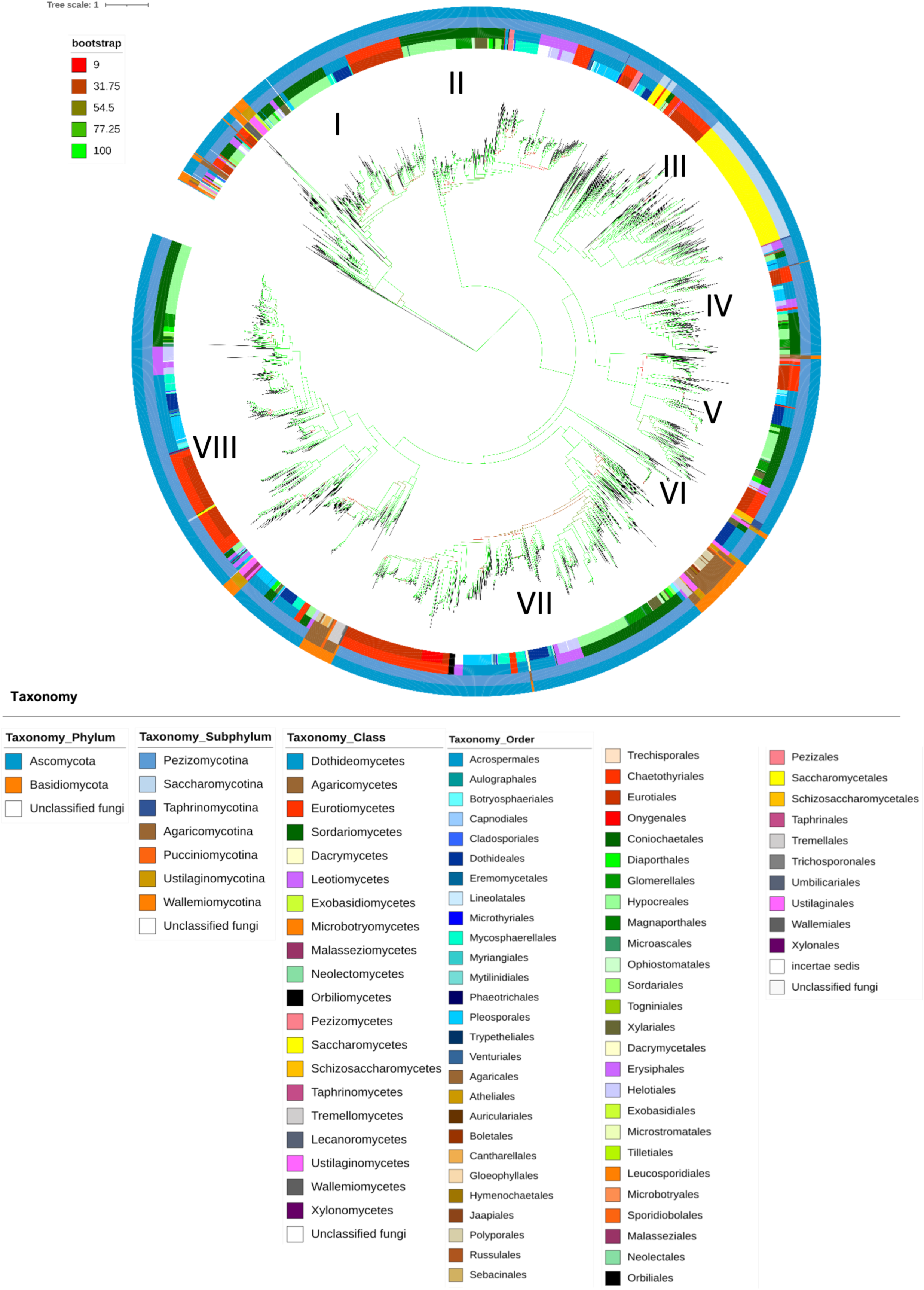
Maximum likelihood phylogenetic tree of fungal acid phosphatase and phytase proteins. This analysis compares the phylogenic relationships among 3058 fungal amino acid sequences annotated as “acid phosphatase” or “phytase” in the UniProtKB database. Detailed metadata for proteins sequences is provided in Table S2. The tree was constructed using the maximum likelihood method with the WAG+I+G4 model of amino acid sequence evolution in IQ-TREE. Branch colors represent confidence intervals based on bootstrapp analyses with 1000 replicates, as indicated in the figure legend. Taxonomic labels were added to visualize the taxonomic distribution with concentric circles representing different taxonomic levels, from phylum (outer ring) to order (inner ring).

**Table 1.**
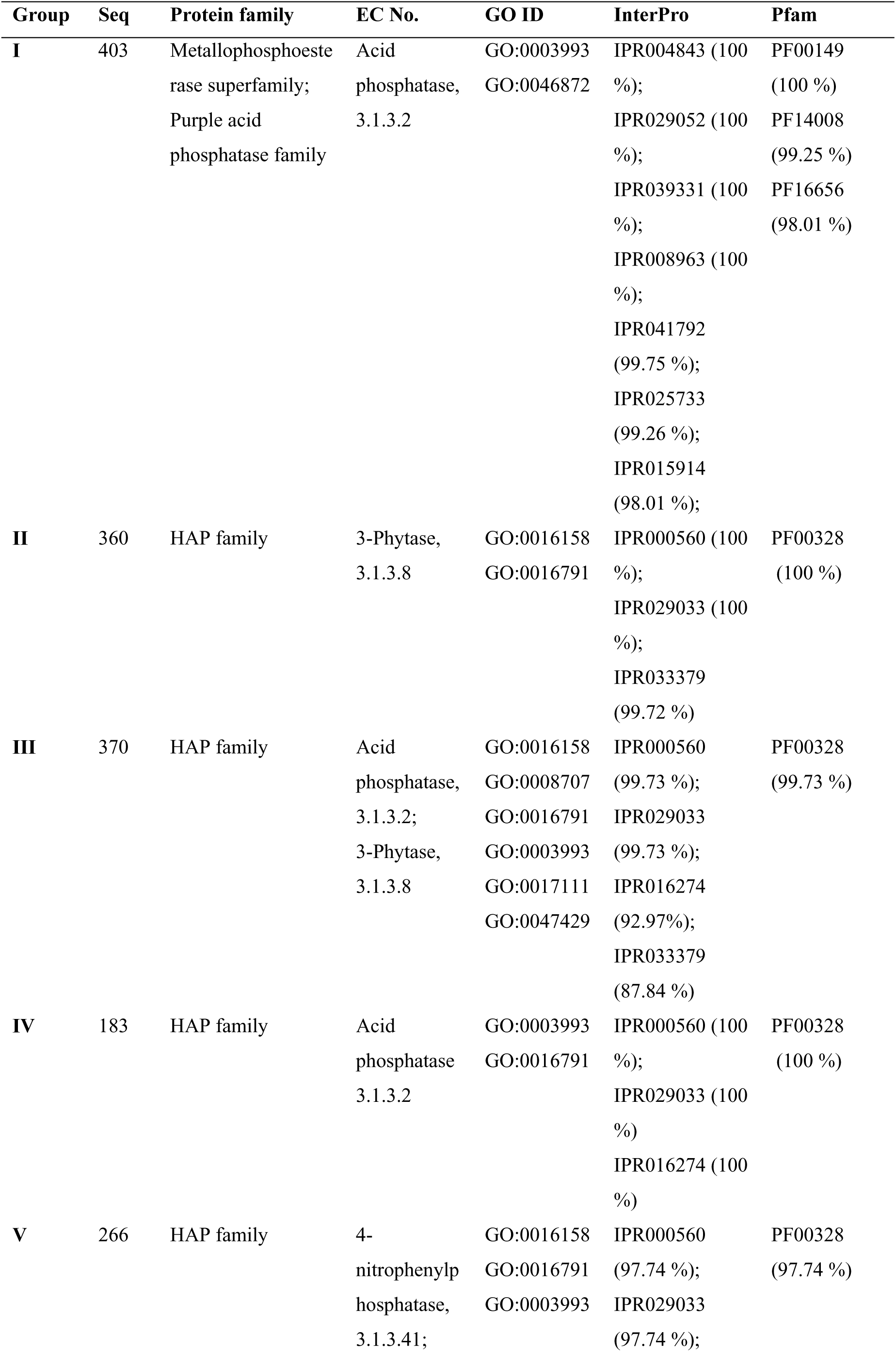
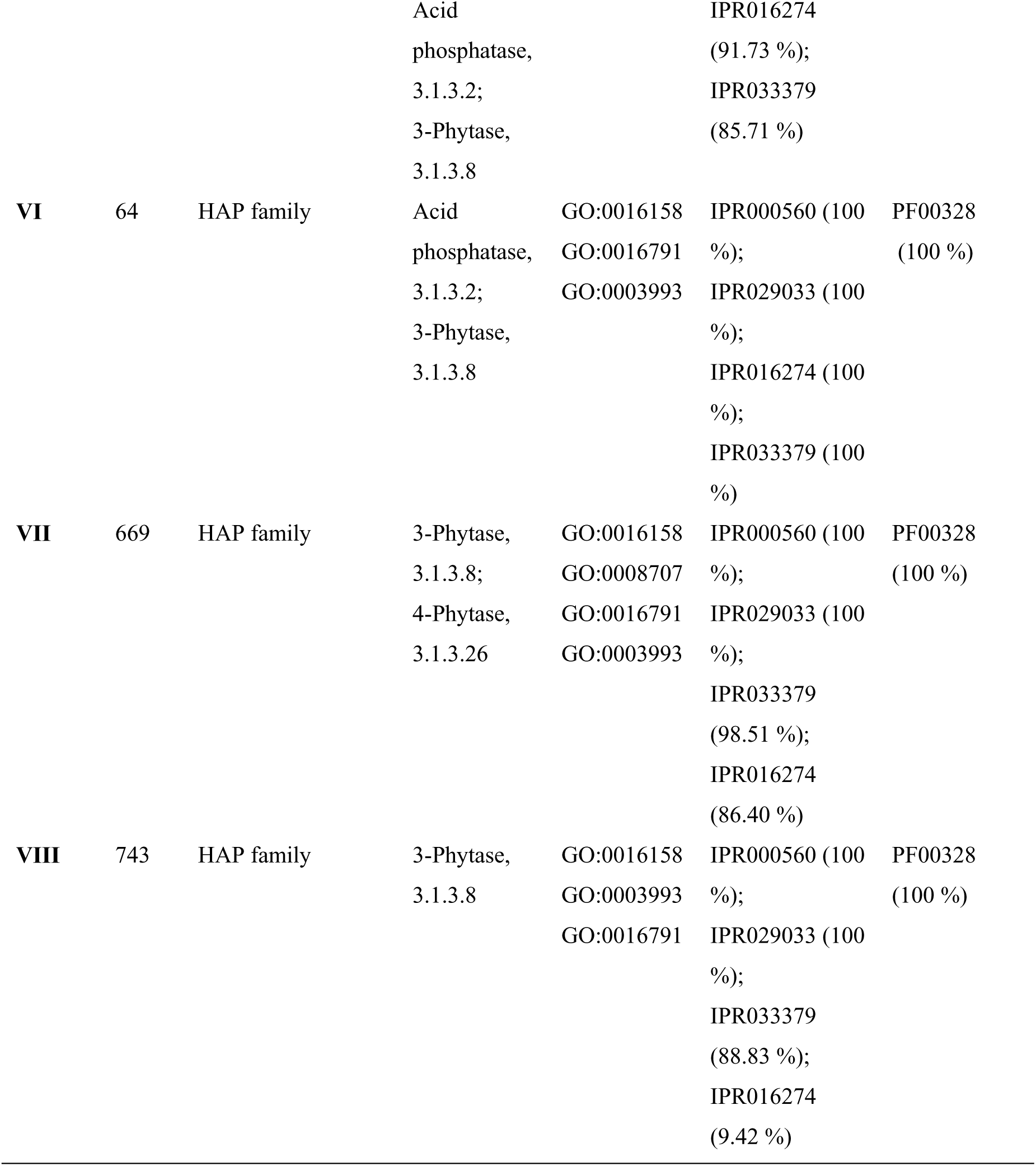
Detailed information of major groups of fungal phosphatase sequences identified in the phylogenetic tree. Cross-references of annotated domains available in UniprotKB from InterPro and Pfam databases are also provided with entry codes exclusive to a group highlighted in bold; their percentage of occurrence in each group is indicated in parentheses. Gene Ontology (GO) IDs, molecular function: acid phosphatase activity, GO:0003993; metal ion binding, GO:0046872; phosphatase activity, GO:0016791; nucleoside-triphosphate diphosphatase activity, GO:0047429; nucleoside-triphosphatase activity, GO:0017111; 3-phytase activity, GO:0016158; 4-phytase activity, GO:0008707; InterPro codes: Calcineurin-like phosphoesterase domain, ApaH type, IPR004843; metallo-dependent phosphatase-like, IPR029052; purple acid phosphatase-like, IPR039331; purple acid phosphatase-like N-terminal, IPR008963; purple acid phosphatase, metallophosphatase domain, IPR041792; iron/zinc purple acid phosphatase-like C-terminal domain, IPR025733; purple acid phosphatase, N-terminal, IPR015914; histidine phosphatase superfamily, clade-2, IPR000560; histidine phosphatase superfamily, IPR029033; HAP active site, IPR033379; HAP, eukaryotic, IPR016274. Pfam codes: calcineurin-like phosphoesterase, PF00149; iron/zinc purple acid phosphatase-like protein C, PF14008; purple acid Phosphatase, N-terminal domain, PF16656; histidine phosphatase superfamily (branch 2), PF00328. Extended information on protein sequences is provided in Table S2.

Fungal phosphatase sequences identified in the phylogenetic tree included proteins annotated as fungal purple acid phosphatases (PAPs) (Group I), acid phosphatases (Group IV), phytases (Groups II, VII and VIII), and groups containing both phytases and acid phosphatases (Groups III, V, and VI). Remarkably, most phytase-containing groups included sequences annotated with Gene Ontology (GO) term GO:0016158, which corresponds to enzymes with 3-phytase activity. Additionally, Groups III and VII also presented sequences annotated with GO:0008707, characteristic of enzymes with 4-phytase activity.

We found that Group I, the most evolutionarily distinct group, comprised 403 protein sequences (13.18 % of the dataset), all annotated as PAPs. These enzymes belong to the metallophosphoesterase superfamily, associated with EC 3.1.3.2. and GO terms GO:0003993 or GO:0046872. The distinct clustering of Group I sequences observed in the phylogenetic tree reflects the early divergence of PAPs from the others. This group was predominantly represented by enzymes from *Aspergillus*, *Talaromyces*, and *Trichoderma*, which are known fungi for their roles in nutrient mobilization and phosphate solubilization in low-P environments, as well as for their ability to colonize plant and animal hosts. Group II includes 360 fungal protein sequences (11.77 % of the total number of sequences). These sequences were mostly annotated as 3-phytases within the HAP family, associated with EC 3.1.3.8. and GO terms GO:0016158 and GO:0016791. This group was taxonomically diverse but especially enriched in the genera *Colletotrichum*, *Fusarium*, and *Verticillium*, many of which are plant-associated, including phytopathogens and endophytes. Group III consisted of 370 sequences (12.10 %), annotated as either acid phosphatase or phytase. These belong to the HAP family, with EC 3.1.3.2. or 3.1.3.8, and GO terms GO:0016158, GO:0008707, GO:0016791 or GO:0003993. This group had broad functional annotation and included members of Saccharomycetales such as *Candida*, *Pichia*, and *Saccharomyces*, suggesting a strong representation of yeasts, many of which are associated with human hosts, fermentation, or opportunistic pathogens. Group IV includes183 sequences, annotated solely as acid phosphatases (5.98 % of the total number of sequences), within the HAP family, EC 3.1.3.2, and with GO terms GO:0016791 or GO:0003993. This group was taxonomically diverse, including filamentous fungi and yeasts such as *Kluyveromyces*, *Candida* and *Yarrowia,-* spanning both commensal yeast and saprotrophic fungi. Group V comprises 266 sequences (8.70 % of the total number of sequences), represented a group containing enzymes annotated a putative acid phosphatase and phytase within the HAP family, with EC 3.1.3.2. or 3.1.3.8 and GO terms GO:0016158, GO:0016791 or GO:0003993. Members of this group included representatives from *Beauveria*, *Trichoderma*, and *Thermomyces*, indicating prevalence of fungi with roles in entomopathogenic activity and thermotolerance. Group VI, the smallest group with 64 protein sequences (2.09 %), also contained proteins annotated as either acid phosphatase or phytase within the HAP family, with EC 3.1.3.8. or 3.1.3.2 and with any of the following GO terms: GO:0016158, GO:0016791 or GO:0003993. Organisms in this group are mostly filamentous saprotrophs with thermotolerant or stress-resistant traits, such as *Thermoascus*, *Myceliophthora*, and *Talaromyces,* known for their thermophilic or stress-resistant traits and roles in cellulose degradation and composting (Alvarez et al, 2016). Group VII contained 669 protein sequences (21,88 %), annotated mainly as phytases from the HAP family, with EC 3.1.3.8 or 3.1.3.26 and GO terms: GO:0016158, GO:0008707, GO:0016791 or GO:0003993. This group was taxonomically broad, including thermophilic and soil fungi such as *Rasamsonia*, *Rhizomucor*, and *Myceliophthora*, often found in compost and decaying plant material. Finally, Group VIII, the largest group, comprised 743 protein sequences, all within the HAP family (24.30 %), and associated with EC 3.1.3.8 and GO terms GO:0016158 or GO:0016791 or GO:0003993. This group encompassed a broad diversity of fungi across various lifestyles, including model organisms, opportunistic pathogens, and symbionts such as *Neurospora*, *Candida*, *Aspergillus*, and *Pichia*, underscoring its functional and evolutionary versatility. Together, these eight groups underscore the evolutionary complexity and functional diversity of fungal acid phosphatases, reflecting diverse ecological strategies and environmental adaptations.

The proteins analyzed were widely distributed across the Fungal Kingdom, spanning two phyla, seven subphyla, 20 classes, 63 orders and 175 families. The dataset spans numerous fungal taxa and ecological strategies, ranging from saprophytes to pathogens and symbionts, emphasizing the key role of acid phosphatases in phosphorus cycling and fungal adaptation to environmental constraints (Fig. 1, Fig. 2 and Table S2). However, the dataset was dominated by sequences from *Ascomycota*, which represented approximately 92% of those used to construct the phylogenetic tree, while only 8% originated from *Basidiomycota*. While Ascomycetes include many soil saprotrophs, endophytes, and plant-associated fungi that often thrive in nutrient-poor environments where phosphate is limiting, this imbalance may reflect biases in available sequencing data, influenced by economic interests, relevance to human or plant pathogens, use as model organisms, or limitations in cultivation. Among *Ascomycota*, the order *Eurotiales* was the most represented, contributing 580 protein sequences. Within this group the most frequent genera were *Aspergillus* (71%), *Penicillium* (17%) and *Talaromyces* (5.2%), while other fungal genera such as *Penicilliopsis* and *Thermomyces* were sparsely represented. Additionally, other highly represented fungal orders included *Hypocreales* (508 proteins, e.g. *Trichoderma* spp., *Fusarium* spp., *Beauveria* spp., *Cordyceps* spp. *Metarhizium* spp.), Pleosporales (234 proteins, e.g. *Alternaria* spp., *Cochliobolus* spp., *Pyrenophora* spp., *Ascochyta* spp.) and *Saccharomycetales* (233 proteins, e.g. *Candida* spp., *Saccharomyces* spp., *Lachancea* spp., *Kluyveromyces* spp., *Pichia* spp., *Hanseniaspora* spp.).

**Figure 2.**
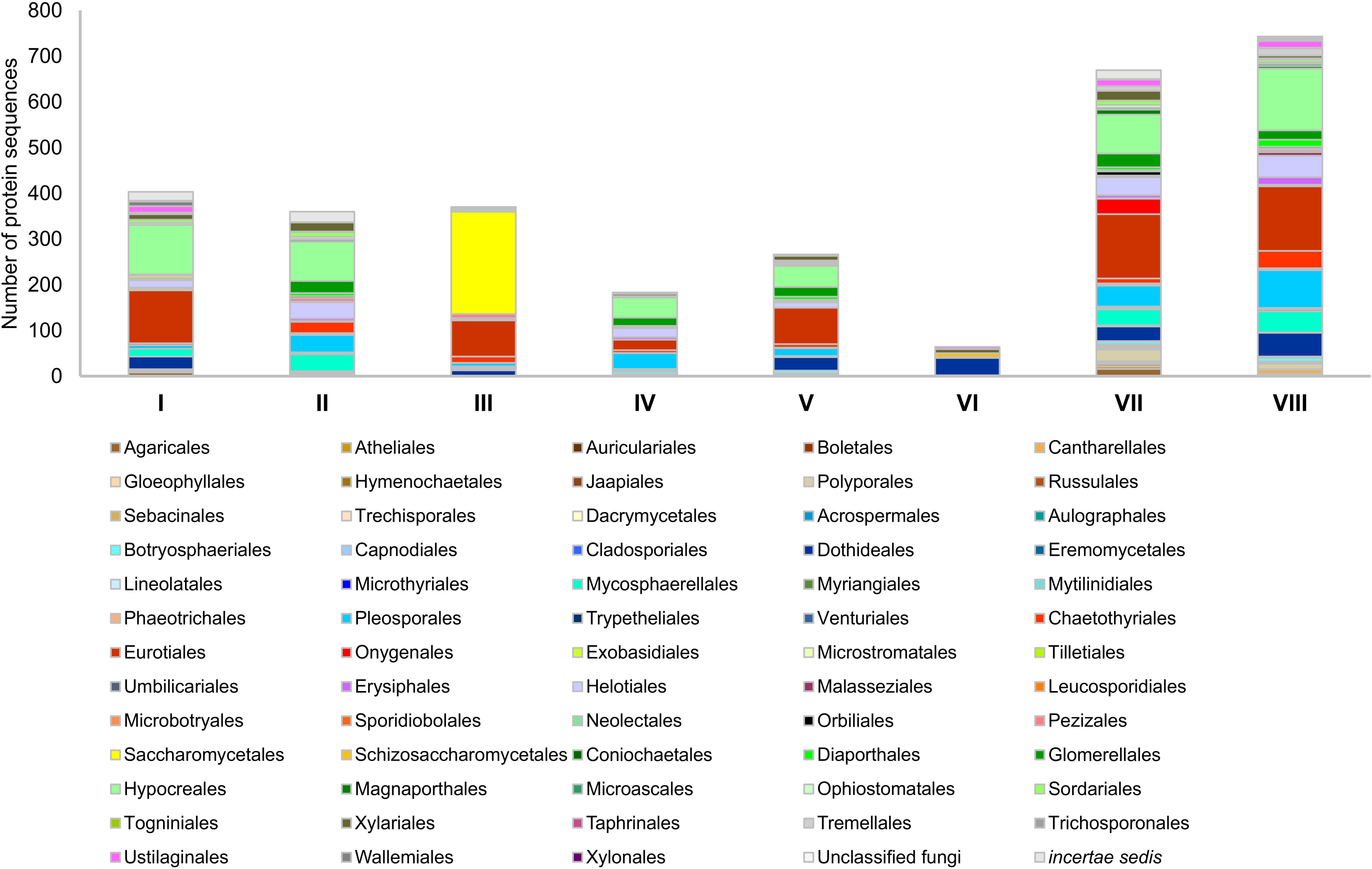
Taxonomic distribution protein sequences in each phosphatase group. The figure shows the number of sequences per group at the order level, based on the dataset used to construct the phylogenetic tree. Taxonomic groups were assigned according to UniProtKB annotations.

All eight groups of fungal phosphatases were found within *Ascomycota*, although with varying distribution. In *Eurotiales*, *Hypocreales* and *Pleosporales*, all the groups were identified except Group VI. In contrast, 96% of *Saccharomycetales* sequences were grouped in Group III, while other groups (I and VIII) were minimally represented, and Groups II, IV, V and VI and VII were absent (Fig. 2).

Within *Basidiomycota*, the most abundant orders were *Ustilaginales* (e.g. *Ustilago* spp., *Sporisorium* spp., *Pseudozyma* spp.) with 48 protein sequences (20%), followed by Polyporales (e.g. *Trametes* spp., *Sparassis* spp., *Dichomitus* spp.) with 42 proteins (18 %), Agaricales (e.g. *Amanita* spp*., Armillaria* spp., *Laccaria* spp., *Leucoagaricus* spp.) with 33 protein sequences (14%) and Tremellales (*Cryptococcus* spp., *Kwoniella* spp., *Naematelia* spp.) with 30 protein sequences (13 %). Other orders, such as *Wallemiales*, *Malasseziales*, and *Boletales* were less represented, comprising 0.4-6% of *Basidimycota*-derived sequences. Interestingly, within *Basidiomycota,* no sequences were assigned to Groups II and III, and Groups IV, V and VI were poorly represented. The majority of sequences clustered in Group VII (41%), VIII (32%) and I (21%) (Fig. 2). Although these data are shaped by the scope of information available in public databases, they lay the foundation for future studies. These sets of results support the hypothesis that constructing fungal acid phosphatase profiles and applying them to metagenomic datasets may help to discover new protein sequences with currently unknown phosphatase activity, enhancing our knowledge of their roles in P-cycling within ecosystems.

### Construction and validation of PROSITE Generalized Profiles for fungal acid phosphatases

To date, no protein profiles specific to fungal phosphatases have been described. As a first step toward distinguishing the different groups identified in the phylogenetic tree, we explored the construction of PROSITE generalized profiles, which are not available in the PROSITE database (https://prosite.expasy.org/) as previously done for bacteria by Udaondo *et al*. (2020).

The construction of protein profiles involves converting multiple sequence alignments into weighted matrices that assign numerical values to matches, mismatches insertions and deletions at each position. These values are summed and compared against a threshold (Sigrist *et al*., 2002) to determine significance. Z-scores equal to or greater than 8.5 are commonly considered biologically meaningful for assigning a protein sequence to a profile (Sigrist *et al*., 2002; Gallegos et al., 1997; Godoy *et al*., 2010; Udaondo *et al*., 2020); although higher thresholds are sometimes used to enhance sensitivity (e.g. Gallegos *et al*., 1997).

PROSITE generalized profiles were constructed using full-length sequences representative of each of the eight fungal phosphatase groups identified in the phylogenetic tree. Following the procedures described in Materials and Methods, one profile was generated for each group. These analyses led to the development of distinct and specific profiles for Group I (referred to as Pfr-A-Fungal_phos) and Group II (referred to as Pfr-B-Fungal_phos), each of which exclusively recognized sequences belonging to its respective group. In contrast, the profiles generated for Groups III to VIII lacked sufficient discriminatory power to distinguish these groups individually. However, none of these profiles cross-recognize protein sequences from Groups I and II. This observation aligns with the phylogenetic tree structure, which shows that proteins from Groups III to VIII are more closely related to each other than to those of Groups I and II. The greater sequence homology among Groups III to VIII, likely accounts for the limited group-level resolution in their profiles (Fig. 3). Thus, we were prompted to construct a broader profile encompassing sequences from Groups III to VIII, which was developed and referred to as Pfr-C-Fungal_phos. The construction of a combined profile for these six groups was further supported by heatmap analyses based on the clustering of the sequences of the study by sequence similarity, which showed clustering of proteins from Groups III to VIII apart from those in Groups I and II (Fig. 3).

**Figure 3.**
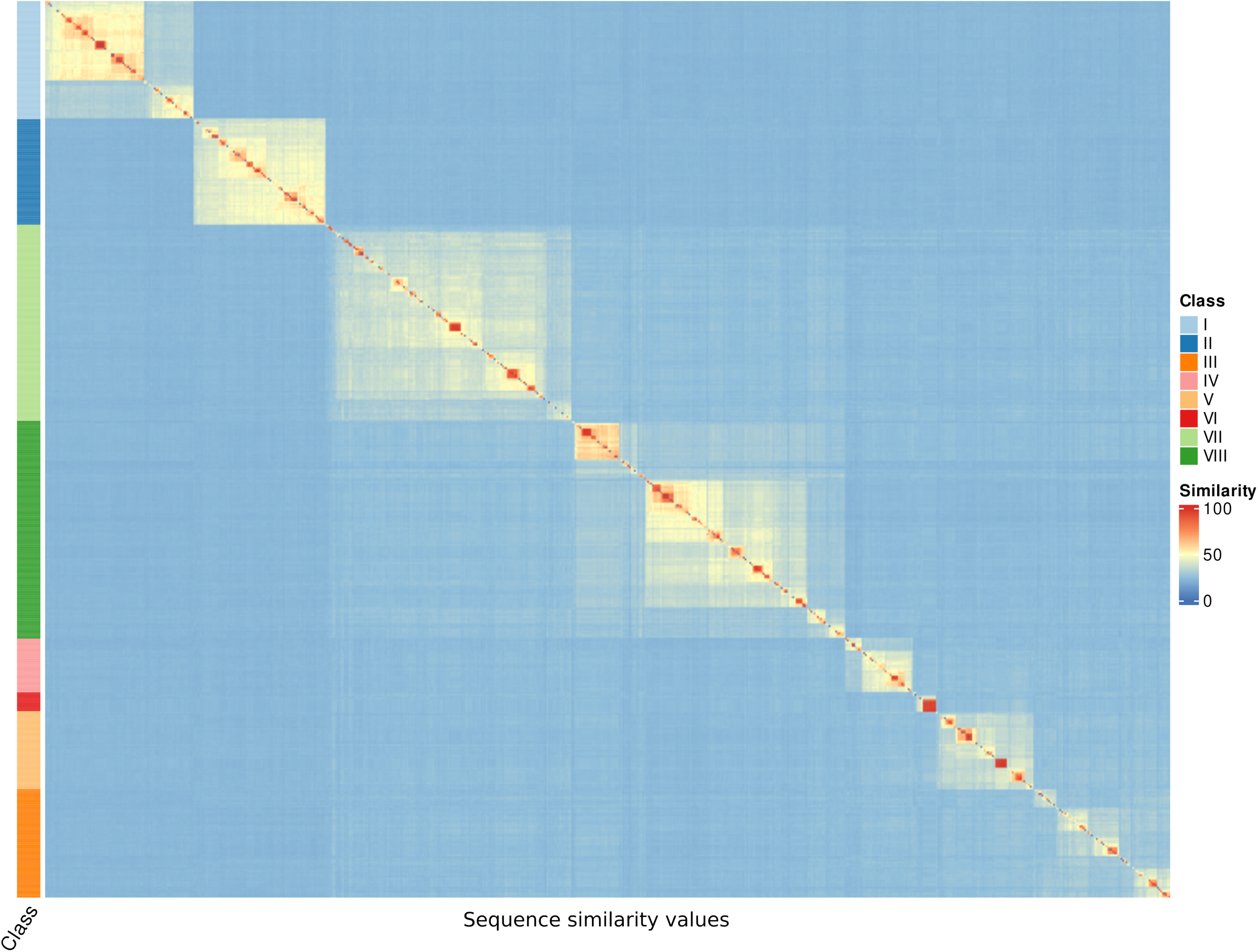
Heat map of sequence similarity among fungal phosphatases. Each colored cell represents the degree of sequence similarity between protein pairs, illustrating clustering patterns that support protein profile definition: Prf-A-Fungal_phos (Group I), Prf-B-Fungal_phos (Group II) and Prf-C-Fungal_phos (Groups from III to VIII) .

When applied to the complete set of 3,058 protein sequences from the phylogenetic tree, Pfr-A-Fungal_phos identified all 403 Group I proteins, with Z-scores ranging from 21.70 to 80.04 (Table 2 and Table S3); Prf-B-Fungal_phos retrieved 360 Group II proteins, with Z-score values ranging from 51.83 to 73.56 (Table 2 and Table S4); and finally, Prf-C-Fungal_phos, which includes more heterogeneous sequences, retrieved 2,295 proteins from Groups III to VIII, with Z-scores ranging from 10.43 to 33.43, matching the total number of proteins in these groups (Table 2 and Table S5). Notably, no false positives (i.e. sequences incorrectly matched to a profile) or false negatives (i.e. sequences that should have matched but were not detected) were observed. This confirms the high specificity of all three profiles, which together comprehensively covered all sequences used in the phylogenetic tree (Table 2).

**Table 2.** Number of sequences recognized by each of the constructed protein profiles for fungal phosphatases within each group of sequences displayed in the phylogenetic tree. The range of Z-score values of sequences retrieved is also indicated. Raw information of the list of hits found by each profile including detailed Z-score value is provided in Table S3 (A, Prf-A-Fungal_phos); Table S4 (B, Prf-B-Fungal_phos) and Table S5 (C, Prf-C-Fungal_phos).

| Profile | I | II | III | IV | V | VI | VII | VIII | Total | Z-score<br>(min-max) |
| --- | --- | --- | --- | --- | --- | --- | --- | --- | --- | --- |
| **A** | 403 | 0 | 0 | 0 | 0 | 0 | 0 | 0 | 403 | 21.699 – 80.035 |
| **B** | 0 | 360 | 0 | 0 | 0 | 0 | 0 | 0 | 360 | 51.830 – 73.564 |
| **C** | 0 | 0 | 370 | 183 | 266 | 64 | 669 | 743 | 2,295 | 10.426 – 33.427 |

In order to further assess their sensitivity and validate the profiles, each was tested against larger UniProtKB datasets as detailed in Material and Methods. We found that the number of sequences retrieved increased for each profile, yet specificity and discriminatory power were maintained, even for those with lower Z-scores, which supports the robustness of the three profiles (Fig. 4; Table S3 to Table S5). The higher number of hits in the UniProtKB datasets was attributed to their ability to detect incomplete or non-annotated sequences that nonetheless contained the hallmark domains of the corresponding groups (Table S3 to Table S5).

**Figure 4.**
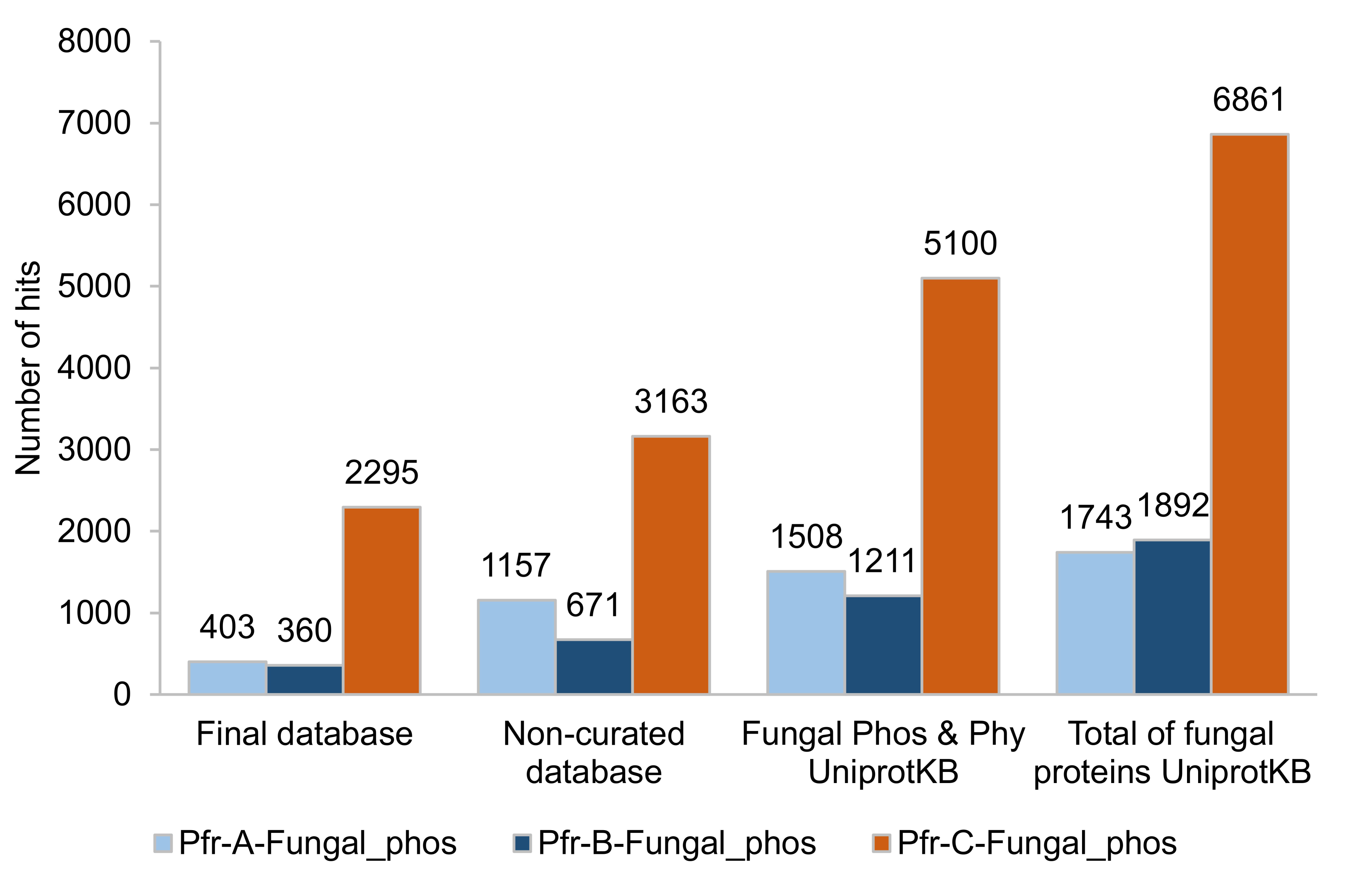
Number of hits detected by each of the three constructed profiles (Prf-A-Fungal_phos; Prf-B-Fungal_phos; Prf-C-Fungal_phos) for fungal phosphatases in different UniProtKB datasets. The datasets include: “Final database”, the 3,058 protein sequences used to construct the phylogenetic tree. “Non-curated database”, 8,859 non-reviewed fungal phosphatase sequences; “Fungal Phos & Phy UniProtKB”, 20,050 fungal sequences annotated as “acid phosphatase” or “phytase” in UniProtKB database and “Total of fungal proteins UniProtKB”, including the complete collection of fungal protein sequences in UniProtKB database (19,274,552 fungal sequences). Detailed protein entries and Z-score for each profile in the corresponding datasets are provided in Table S3 (Prf-A-Fungal_phos), Table S4 (Prf-B-Fungal_phos) and Table S5 (Prf-C-Fungal_phos).

Additionally, the three fungal phosphatase profiles were used to search a filtered UniRef 90 database comprising 29,638,836 clusters of proteins annotated as hypothetical, unknown or uncharacterized proteins. The Pfr-A-Fungal_phos profile identified 44 uncharacterized protein sequences across a broad taxonomic spectrum with Z-scores ranging from 8.50 to 77.73. These sequences were taxonomically diverse, spanning both *Ascomycota* and *Basidiomycota*. Within *Ascomycota*, many hits were affiliated with genera such as *Penicillium*, *Aspergillus*, *Apiospora*, and *Rhizomucor*, which are known for their roles in soil, plant association, or industrial applications. *Penicillium* was particularly well represented, accounting for more than one-third of the hits. Sequences were also retrieved from less commonly studied genera such as *Recurvomyces*, *Ophidiomyces*, and *Priceomyce*. In *Basidiomycota*, the profile retrieved sequences from *Amanita muscaria* and *Stereum hirsutum*, highlighting the ability of the Pfr-A-Fungal_phos profile to detect distantly related phosphatases. The taxonomic distribution of the uncharacterized proteins retrieved with this profile is shown in Figure 5 and Suppl. Table 6. The Pfr-B-Fungal_phos profile retrieved 104 fungal protein sequences from the UniRef90 database, annotated as uncharacterized, hypothetical, or unknown proteins with Z-scores ranging from 8.51 to 70.41. We were able to identify candidate phosphatases sequences from both major fungal phyla, *Ascomycota* and *Basidiomycota*, as well as from early-diverging lineages such as *Rozellomycota*, *Mucoromycota*, and *Zoopagomycota*. Notably, sequences were identified in species from diverse ecological niches, including plant pathogens (e.g., *Rhizoctonia solani*), symbionts (e.g., *Rozella allomycis*), endophytes, and thermophilic fungi (Fig. 6 and Suppl. Table 7). The Pfr-C-Fungal_phos profile retrieved a total of 293 fungal protein sequences from the UniRef90 database, all annotated as hypothetical, uncharacterized, or unknown proteins, with Z-scores ranging from 8.53 to 19.63. This profile captured the broadest taxonomic range among the three, reflecting its design to detect homologs across phosphatase Groups III to VIII. The majority of sequences (over 80%) belonged to the phylum *Ascomycota*, with *Penicillium* once again standing out as the most represented genus. Other frequently identified genera included *Aspergillus*, *Talaromyces*, *Fusarium*, and *Trichoderma*, all known for their ecological and industrial significance. Within *Basidiomycota*, the profile successfully detected proteins from species such as *Schizophyllum commune*, *Fomitopsis pinicola*, and *Rhizoctonia solani*, highlighting its utility in recognizing acid phosphatase homologs in both saprophytic and pathogenic lineages (Fig. 7 and Suppl. Table 8).

**Figure 5.**
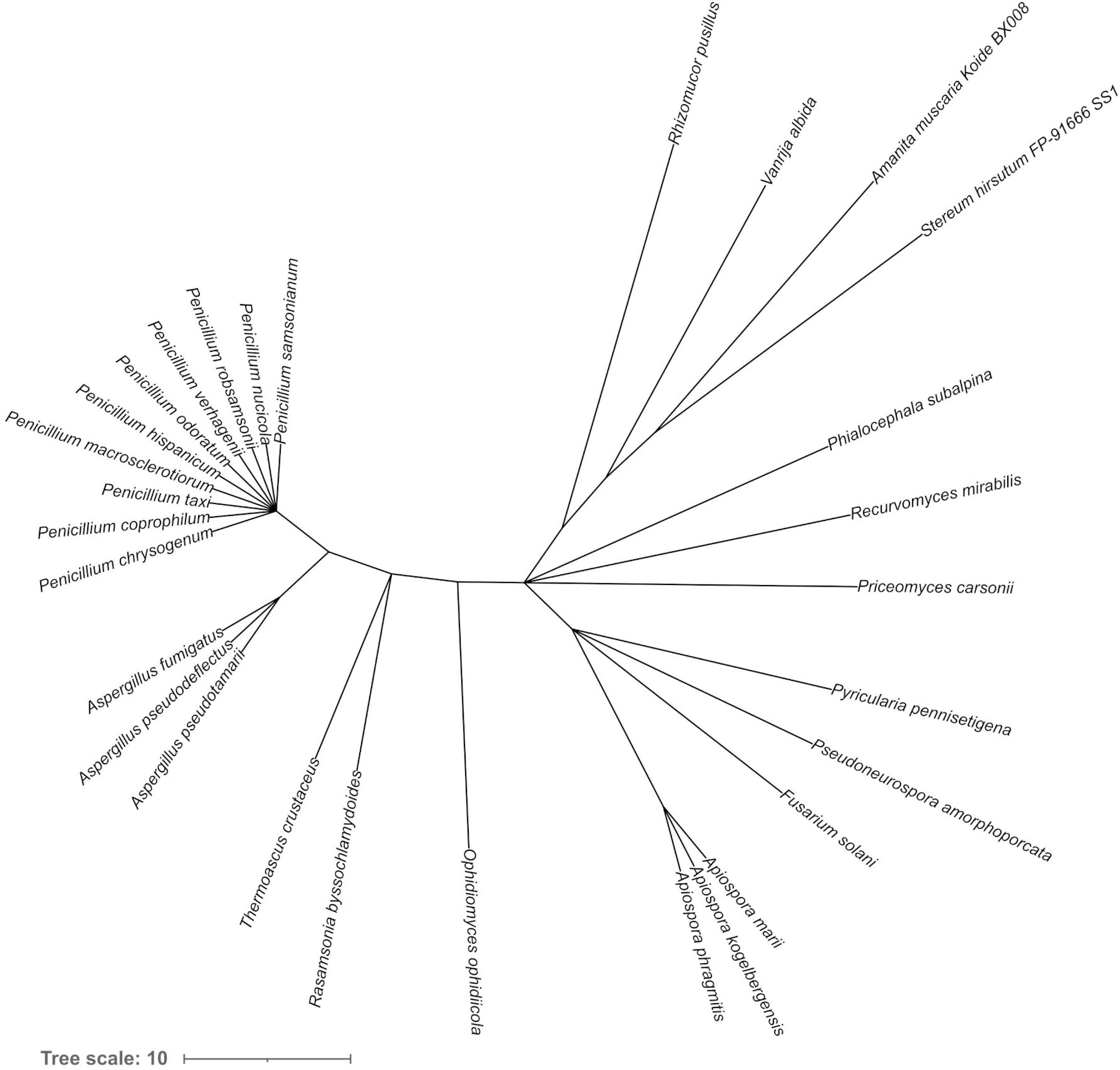
Taxonomic distribution of proteins annotated as “hypothetical protein”, “unknown protein” or “uncharacterized protein” from UniRef90 and retrieved using acid phosphatase profiles. Taxonomic identifiers from the matched sequences were mapped and visualized using the Taxonomy Browser from the NCBI, to illustrate the phylogenetic diversity of the candidate phosphatases identified by the profiles Prf-A-Fungal_phos,

**Figure 6.**
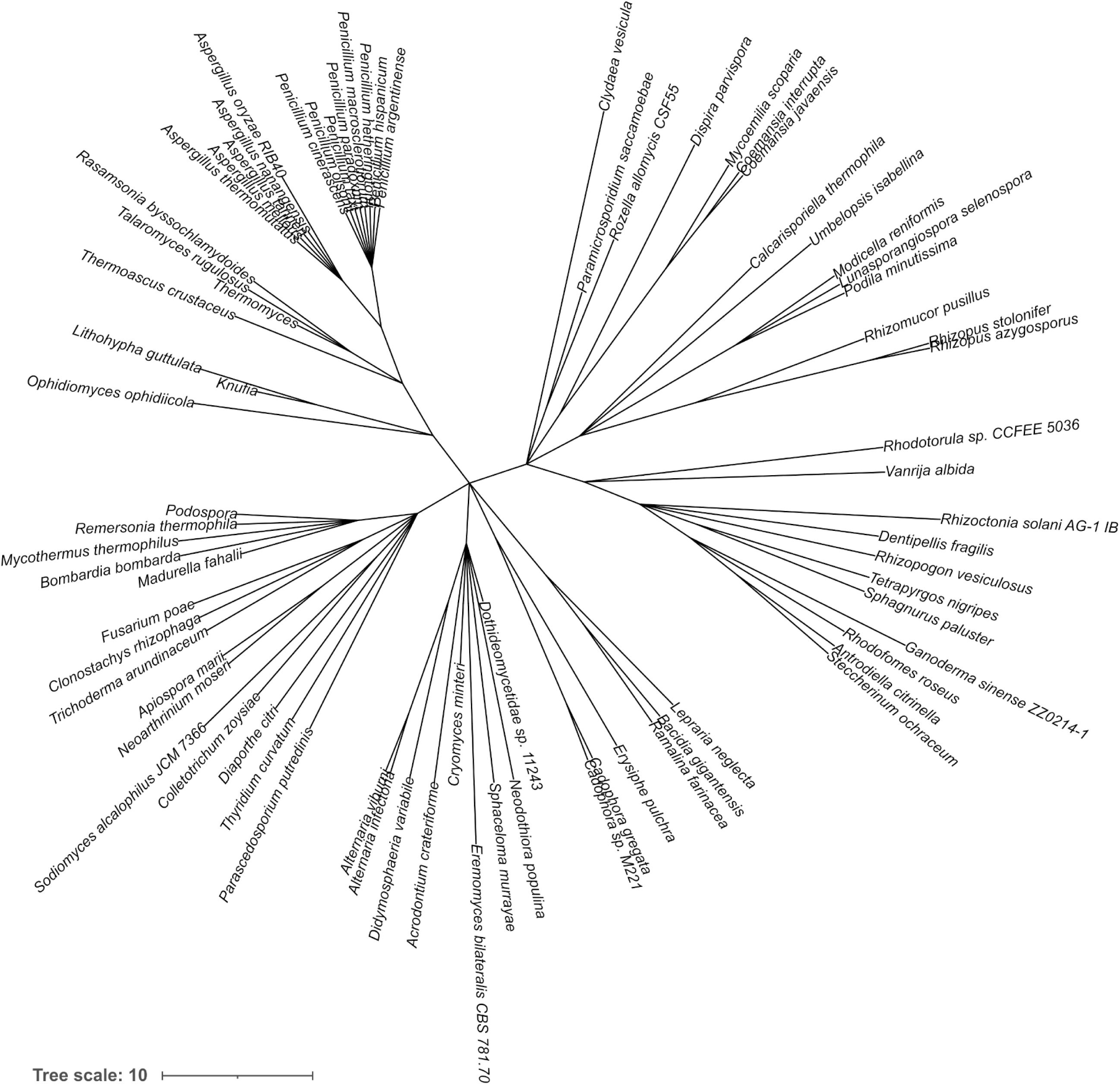
Taxonomic distribution of proteins annotated as “hypothetical protein”, “unknown protein” or “uncharacterized protein” from UniRef90 and retrieved using acid phosphatase profiles. Taxonomic identifiers from the matched sequences were mapped and visualized using the Taxonomy Browser from the NCBI, to illustrate the phylogenetic diversity of the candidate phosphatases identified by the profile Prf-B-Fungal_phos

**Figure 7.**
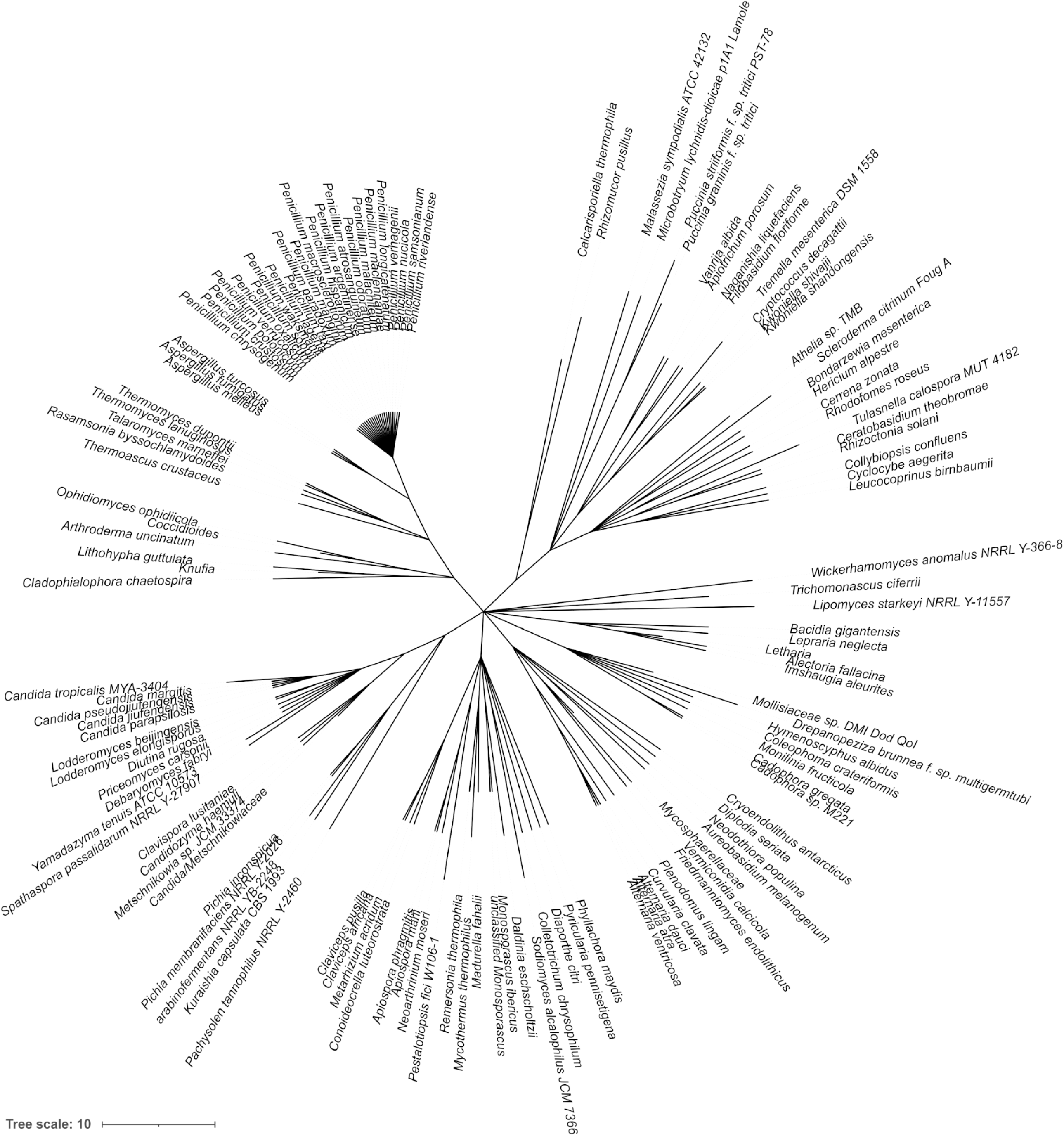
Taxonomic distribution of proteins annotated as “hypothetical protein”, “unknown protein” or “uncharacterized protein” from UniRef90 and retrieved using acid phosphatase profiles. Taxonomic identifiers from the matched sequences were mapped and visualized using the Taxonomy Browser from the NCBI, to illustrate the phylogenetic diversity of the candidate phosphatases identified by the profile Prf-C-Fungal_phos.

These results highlight the potential utility of the designed fungal acid phosphatase profiles for uncovering a previously uncharacterized and widespread phosphatase family in fungi. The wide taxonomic distribution of the lineages captures for the three profiles demonstrates their capacity to detect distantly related and potentially novel acid phosphatase candidates from across the fungal tree of life. The diversity of identified taxa, spanning from industrial workhorses to cryptic soil fungi and obligate symbionts, emphasizes the broad functional relevance of acid phosphatases.

### Experimental validation and ecological implications of fungal phosphatase profiles

As a proof of concept to empirically validate these profiles, we selected one non-reviewed protein sequence from each profile with a high Z-score. The DNA sequences were synthesized via reverse genetics, cloned into the pRD196 expression vector, and expressed in *S. cerevisiae* strain BY4741. Cells were grown to the late exponential phase under constitutive expression of the respective proteins. The cells were harvested and cell-free extracts prepared as described in Materials and Methods, and both acid phosphatase and phytase activities were assessed. The results presented in Suppl. Table 9 confirmed that all three proteins exhibited acid phosphatase activity at pH 5.5 with pNPP (3.6 to 7.6 3 U/mg protein) as a substrate. This validates the functional annotation of previously uncharacterized sequences as acid phosphatases. No phytase activity was detected within the limited set of enzymes tested.

In summary, the phylogenetic tree constructed, encompassing over 3,000 proteins, provides unequivocal insights into the complexity of fungal acid phosphatases as crucial enzymes involved in the P cycle, particularly in the release of inorganic phosphate from organophosphorus molecules. The fungal phosphatase profiles developed in this study, together with existing bacterial profiles (Udaondo *et al*., 2020), represent powerful tools to explore the full diversity of these key enzymes in both soil and aquatic environments. These profiles not only pave the way for the discovery of novel phosphatases, but also contribute to a deeper understanding of their roles in environmental phosphorous cycling, opening new avenues for biotechnological applications in phosphorous recovery, particularly in nutrient-poor soils and contaminated environments. As the demand for sustainable agricultural practices and efficient nutrient management grows, the development and application of novel, and biotechnologically improved acid phosphatases, could play a critical role in improving soil fertility and promoting environmental sustainability, ultimately contributing to the mitigation of global challenges like climate change and resources depletion.

## DATA AVAILABILITY

Protein profiles designed in this study were deposited in the Zenodo database and can be accessed with the entry https://doi.org/10.5281/zenodo.16876021. In addition, datasets analyzed during the present study are provided as supplementary material.

## ACKNOWLEDGMENTS

This work was financially supported by the European Union H2020 Research and Innovation Program under grant agreement No. 862695 (European Joint Program SOIL, TRACE-Soils project). The work in our laboratory was also funded by grants from the Spanish Ministry of Science and Innovation (Plan Nacional PDI-2021-123469OB-I00 and Plan Nacional PID2024-161463NA-I00) and Junta de Andalucía Grant P20-00049 by the Regional Government of Andalusia. Bioinformatic analyses presented here were carried out using the High-Performance Computing resources provided by the Supercomputing and Bioinnovation Center of the University of Malaga, Spain. We thank Angela Tate for critical reading of the manuscript.

## AUTHOR CONTRIBUTIONS

Tamara Gómez-Gallego: methodology, investigation, writing first draft, revise final version Zulema Udaondo: methodology, investigation, validation, writing, revise final version, funding acquisition Rocío Palacios-Ferrer: methodology, investigation, revise final versión Luis Díaz-Mártinez: methodology, revise final versión Juan Luis Ramos: conceptualization, methodology, validation, writing, writing final version, funding acquisition

